# TWIST1 methylation by SETD6 selectively antagonizes LINC-PINT expression in Glioblastoma multiforme

**DOI:** 10.1101/2021.11.18.469142

**Authors:** Lee Admoni-Elisha, Michal Feldman, Tzofit Elbaz, Anand Chopra, Guy Shapira, Christopher J Fry, Noam Shomron, Kyle Biggar, Dan Levy

**Author notes:** Correspondence should be addressed to D.L.

## Abstract

Glioblastoma multiforme (GBM) is the most common and aggressive malignant brain tumor among adults, which is characterized by high invasion, migration and proliferation abilities. One important process that contributes to the invasiveness of GBM is the epithelial to mesenchymal transition (EMT). EMT is regulated by a set of defined transcription factors which tightly regulate this process, among them is the basic helix-loop-helix family member, TWIST1. Here we show that TWIST1 is methylated on lysine-33 at chromatin by SETD6, a methyltransferase with expression levels correlating with poor survival in GBM patients. RNA-seq analysis in U251 GBM cells suggested that both SETD6 and TWIST1 regulate cell adhesion and migration processes. We further show that TWIST1 methylation attenuates the expression of the long-non-coding RNA, LINC-PINT, thereby suppressing EMT in GBM. Mechanistically, TWIST1 methylation represses the transcription of LINC-PINT by increasing the occupancy of EZH2 and the catalysis of the repressive H3K27me3 mark at the LINC-PINT locus. Under un-methylated conditions, TWIST1 dissociates from the LINC-PINT locus, allowing the expression of LINC-PINT which leads to increased cell adhesion and decreased cell migration. Together, our findings unravel a new mechanistic dimension for selective expression of LINC-PINT mediated by TWIST1 methylation.

## Introduction

Glioblastoma multiforme (GBM) is the most common and aggressive malignant brain tumor among adults, with median survival of 1-2 years with 7% of 5-years-survival rate (1,2). GBM tumors are highly diffusive, invasive and vascularized and therefore cannot be cured by surgical intervention (3). One important process that contributes to the invasiveness of GBM is the epithelial to mesenchymal transition (EMT). In this process, epithelial cells undergo multiple changes which include loss of their junctions and apical-basal polarity, cytoskeleton reorganization and increased production of extracellular matrix (ECM) components (4,5). These changes result in enhanced motility, invasiveness, and resistance to apoptosis (6). EMT is regulated by a set of defined transcription factors (TFs), including TWIST1, SNAIL, SLUG, and ZEB1/2 (4,6). Indeed, these TFs were found to play a key role in the development and progression of GBM (7-11).

TWIST1 belongs to the bHLH (basic helix-loop-helix) transcription factors. The human TWIST1 is approximately 21 kDa and contains two nuclear localization sequences (12). TWIST1 binds the DNA sequences ^5^’CANNTG^3^’,named E-boxes, through a conserved bHLH domain. This domain is also important for the interactions with other proteins to form homo- and hetero-dimeric complexes (12,13).

In addition to the physiologic role of TWIST1 in embryonic development, organogenesis and angiogenesis (12,14), this transcription factor is also associated with many types of aggressive tumors (12,15). The most critical pathological function of TWIST1 in cancer is facilitating tumor invasion and metastasis by promoting EMT (16). In GBM, TWIST1 is highly expressed in patient tissue specimens (17). TWIST1 was also found to promote invasion in GBM through the upregulation of genes such as SNAI1, MMP2, HGF, and FN1, which associate with adhesion, extracellular matrix, cell motility and locomotion (11).

TWIST1 is regulated by diverse post-translational modifications. Phosphorylation at S68 promotes the heterodimerization with E12, which leads to a pro-invasive phenotype (18) and prevents its ubiquitination-mediated degradation (19). In contrast, AKT1 and AKT2 phosphorylate TWIST1 at S42, a modification promoting TWIST1 degradation (20). Upon DNA damage, RNF8 mediates K63-linked poly-ubiquitination at K38, leading to TWIST1 stabilization and activation (21). In lung cancer, PRMT1 modulates TWIST1 function through methylation at R34. This methylation was shown to be crucial for the repression of epithelial markers and TWIST1 nuclear localization (22). TWIST1 function is also regulated by di-acetylation at K73 and K76 mediated by the acetyltransferase Tip60. This di-acetylation promotes the TWIST1-BRD4 interaction which activates the transcription of WNT5a (23). Up to date, the regulation of TWIST1 activity by lysine methylation has not been reported yet.

Lysine methylation is catalyzed by protein-lysine methyltransferases. While extensive studies were performed on histone proteins, it is now clear that lysine methylation extends far beyond that, with nearly 3000 non-histone sites reported to be methylated in PhosphoSitePlus (24). However, only a small fraction of these methylation events were functionally studied. The SET domain-containing protein 6 (SETD6) is a member of the lysine methyltransferase family. SETD6 was first enzymatically characterized as a regulator of inflammation through the methylation of NF-kB/RelA protein (25). Later studies revealed its role in a variety of cellular processes and signaling pathways such as transcription, WNT signaling, cell cycle, oxidative stress response, hormone receptor signaling and more (26-31).

Long non-coding RNAs (LncRNAs) are a large heterogeneous group of RNA molecules longer than 200 nucleotides, that are not transcribed into functional proteins. Despite not being transcribed, it is now apparent that LncRNAs are functional molecules that regulate diverse cellular processes (32). Long intergenic p53 induced transcript (LINC-PINT) was first characterized as a target of P53 in mouse cell line, which regulates gene expression through interaction with the Polycomb repressive complex 2 (PRC2) (33). Later studies have shown that LINC-PINT inhibits pro-invasive genes and abolishes the invasiveness of cancer cells (34). Accordingly, LINC-PINT was found to be downregulated in multiple types of cancer such as colorectal cancer, lung adenocarcinoma and Glioblastoma (34,35). Recently, a connection between LINC-PINT to EMT was identified. In GBM, LINC-PINT was found to suppress EMT phenotype by blocking the WNT/β-catenin pathway (35), and in laryngeal squamous cell carcinoma it was found to suppress EMT by inhibiting the transcription factor ZEB1 (36).

Here we show that TWIST1 is regulated by lysine methylation. We have found that TWIST1 is targeted for methylation at chromatin by SETD6 and that high expression of SETD6 correlates with poor survival in GBM patients. RNA-seq experiments in U251 SETD6 depleted cells as well as cells stably expressing TWIST1, revealed a significant enrichment in cellular processes linked to extracellular matrix organization and cell adhesion which are involved in the EMT process during tumorigenesis (37). Our data further provide evidence that the methylation of TWIST1 at K33 selectively regulates the expression of LINC-PINT. Methylated TWIST1 binds to the LINC-PINT locus and limits it transcription by increasing the repressive H3K27me3 mark. The occupancy of unmethylated TWIST1 at the LINC-PINT locus is dramatically reduced and correlates with increased expression of LINC-PINT RNA, resulting in augmented cell adhesion and reduced cell migrations, thereby mimicking the phenotypes of over-expressed LINC-PINT in GBM cells.

## Materials and Methods

### Plasmids

For recombinant purification TWIST1, SNAIL and SLUG sequences were amplified by PCR and subcloned into pET-Duet plasmids. TWIST1 mutants were generated using site-directed mutagenesis and cloned into pET-Duet. Primers used for cloning and mutagenesis are listed in **Table1**. TWIST1 WT and K33R were further cloned into pcDNA3.1 3xFLAG. SETD6 was cloned into pcDNA3.1 3xHA plasmid. For viral infection, TWIST1 WT and K33R were cloned into pWZL-FLAG plasmid. pBABE plasmid containing LINC-PINT cDNA (BC130416.1) were kindly provided by Dr. Maite Huarte (University of Navarra, Spain).

**Table 1.**
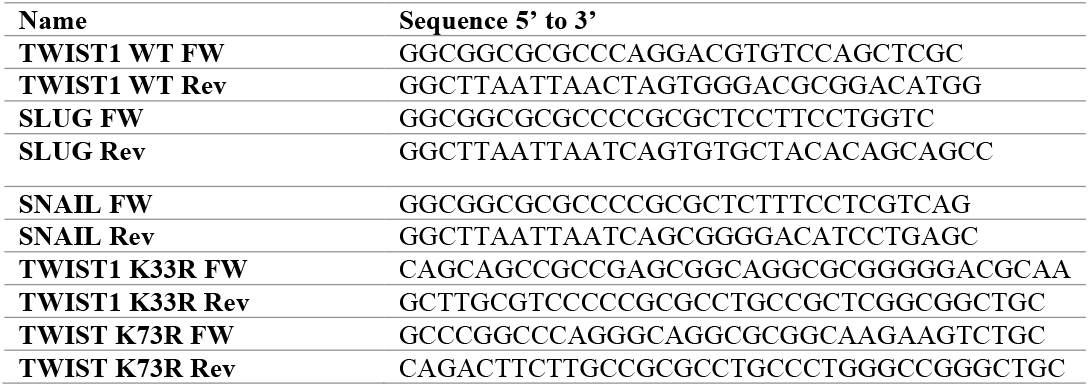
Primers for cloning and mutagenesis

### Cell lines, transfection, infection and treatment

Human Glioblastoma cell lines U251, Human embryonic kidney cells (HEK293T) were maintained in Dulbecco’s modified Eagle’s medium (Sigma, D5671) with 10% fetal bovine serum (FBS) (Gibco), penicillin-streptomycin (Sigma, P0781), 2 mg/ml L-glutamine (Sigma, G7513) and non-essential amino acids (Sigma, M7145), at 37°C in a humidified incubator with 5% CO2.

Cell transfections were performed using polyethyleneimine (PEI) reagent (Polyscience Inc., 23966) or jetPRIME (Polyplus transfection, 114-07) according to manufacturer’s protocol.

For CRISPR/Cas9 SETD6 knock-out, four different gRNAs for SETD6 were cloned into lentiCRISPR plasmid (Addgene, #49535). Following transduction and puromycin selection (2.5μg/ml), single clones were isolated, expanded and validated by sequencing.

For stable transfections in U251 cell line, retroviruses were produced by transfecting HEK293T cells with the indicated pWZL constructs (Empty, FLAG TWIST1 WT, FLAG TWIST1 K33R) or pBABE (LINC-PINT) with plasmids encoding VSV and gag-pol. U251 cells were infected with the viral supernatants and selected with 500 μg/ml hygromycin B (TOKU-E).

### Recombinant Proteins and Peptides

Escherichia coli Rosetta transformed with a plasmid expressing His tagged TWIST1 WT or mutants, SNAIL, SLUG were grown in LB medium. Bacteria were harvested by centrifugation after IPTG induction and lysed by sonication on ice (25% amplitude, 1 min total, 10/5 sec ON/OFF). His-tagged proteins were purified using Ni-NTA beads (Pierce) or on a HisTrap column (GE) with the ÄKTA gel filtration system. Proteins were eluted by 0.5 M imidazole followed by dialysis to 10% glycerol in phosphate-buffered saline (PBS). Recombinant GST SETD6 was expressed and purified as previously described (25).

### Antibodies, Western blot Analysis and Immunoprecipitation

Primary antibodies used were: anti-FLAG (Sigma, F1804), anti-HA (Millipore, 05– 904), anti-Actin (Abcam, ab3280), anti-Pan-methyl (Cell signaling, 14679), anti-GST (Abcam, ab9085), anti-SETD6 (Genetex, GTX629891), anti-TWIST1 (Abcam 50887), anti-His (Thermo Fisher scientific, rd230540a), anti-EZH2 (Cell signaling #5246) anti H3K27me3 (Cell signaling, 9733) and anti-histone3 (H3) (Abcam, ab10799). Anti-TWIST1 K33me1 was generated in collaboration with Cell Signaling Technology. Rabbits were immunized with a synthetic peptide corresponding to residues surrounding mono-methylated Lys33 of human TWIST1 protein. This antibody was purified by peptide affinity chromatography. HRP-conjugated secondary antibodies, goat anti-rabbit, goat anti-mouse, were purchased from Jackson ImmunoResearch (111-035-144, 115-035-062 respectively). For Western blot analysis, cells were homogenized and lysed in RIPA buffer (50 mM Tris-HCl pH 8, 150 mM NaCl, 1% Nonidet P-40, 0.5% sodium deoxycholate, 0.1% SDS, 1 mM DTT, and 1:100 protease inhibitor mixture (Sigma)). Samples were resolved on SDS-PAGE, followed by Western blot analysis. For immunoprecipitation, proteins extracted from cells were incubated overnight at 4°C with FLAG-M2 beads (Sigma, A2220) or pre- conjugated A/G agarose beads (Santa Cruz, SC-2003) with antibody of interest. The beads were then washed three times with RIPA buffer and submitted to SDS-PAGE and Western blot analysis.

### In-Vitro Methylation Assay

Methylation assay reactions contained 1 μg of His-TWIST1 WT or mutant and 4 μg of His SETD6 or GST SETD6, 2 mCi of 3H-labeled S-adenosyl-methionine (SAM) (Perkin-Elmer, AdoMet) and PKMT buffer (20 mM Tris-HCl pH 8, 10% glycerol, 20 mM KCl, 5 mM MgCl2). The reaction tubes were incubated overnight at 30°C. The reactions were resolved by SDS-PAGE for Coomassie staining (Expedeon, InstantBlue) or autoradiography.

For the non-radioactive (cold) methylation assay, 300 μM non-radioactive SAM was added (Abcam, ab142221).

### Semi In-Vitro Methylation Assay

HEK293T cells were transfected with FLAG-TWIST1 WT or K33R plasmids. Chromatin fractions were immunoprecipitated with FLAG-M2 beads overnight at 4°C. The samples were then washed 3 times with dilution buffer and once with PKMT buffer, followed by an *in-vitro* radioactive methylation assay overnight at 30°C, in the presence of 4 μg His-SETD6. The reactions were resolved by SDS-PAGE for Coomassie staining or autoradiography.

### Enzyme-linked Immunosorbent Assay (ELISA)

Approximately 2ug of His-TWIST1, His-SNAIL, His-SLUG or BSA diluted in PBS were added to a 96-well plate (Greiner Microlon) and incubated for 1 hour at room temperature followed by blocking with 3% BSA for 30 min. Then, the plate was covered with 0.5 μg GST-SETD6 or GST protein (negative control) diluted in 1% BSA in PBST for 1 hour at room temperature. Plates were then washed and incubated with primary antibody (anti-GST, 1:4000 dilution) followed by incubation with HRP-conjugated secondary antibody (goat anti-rabbit, 1:2000 dilution) for 1 hour. Finally, TMB reagent and then 1 N H2SO4 were added; the absorbance at 450 nm was detected using Tecan Infinite M200 plate reader.

### Fibronectin Adhesion Assay

For cell adhesion assay to fibronectin, cells were serum starved (0% FBS) overnight. Then, cells were harvested and 1×10^5^ cells/well were plated on a fibronectin (Millipore, 341631) pre-coated 96-well plate (2.5 μg/well) or BSA as a negative control (5% in PBS) for 4 hours, followed by a PBS wash and crystal violet staining (0.5% crystal violet in 20% methanol). Crystal violet staining was solubilized in 2% SDS and quantified at 550 nm using Tecan Infinite M200 plate reader.

### Wound healing migration assay

For migration assay, 1×10^5^ cells were seeded in 24 well plates 1 day before performing the wound. The wound was produced using 200ul pipette tip. Cell migration was monitored for 72 hours, following image processing and wound closure analysis by LionheartTM FX Automated Microscope (4x).

### Transwell migration assay

Serum starved cells (5×10^4^) were seeded onto a semi-permeable membrane inserts (8μm pore size) with 10% FBS medium below. After 24h, inserts were washed with PBS followed by fixation with 3.7% Formaldehyde and permeabilization with 100% Methanol. Inserts were then washed and stained with Geimsa solution (Sigma, 48900). Non-migrated cells were removed using a cotton swab and images were taken using EVOS FL Cell Imaging System (4 fields for each cell type) and analyzed using Fiji software.

### RNA Extraction and Real-Time qPCR

Total RNA was extracted using the NucleoSpin RNA Kit (Macherey-Nagel). 200 ng of the extracted RNA was reverse-transcribed to cDNA using the iScript cDNA Synthesis Kit (Bio-Rad) according to the manufacturer’s instructions. Real-time qPCR was performed using the UPL probe library system or SYBR green I master (Roche) in a LightCycler 480 System (Roche). The real-time qPCR primers were designed using the universal probe library assay design center (Roche) and UCSC Genome browser (**Table 2**). All samples were amplified in triplicates in a 384-well plate using the following cycling conditions: 10 min at 95°C, 45 cycles of 10 sec at 95°C, 30 sec at 60°C and 1 sec at 72°C, followed by 30 sec at 40°C. Gene expression levels were normalized to GAPDH and controls of the experiment.

**Table 2.**
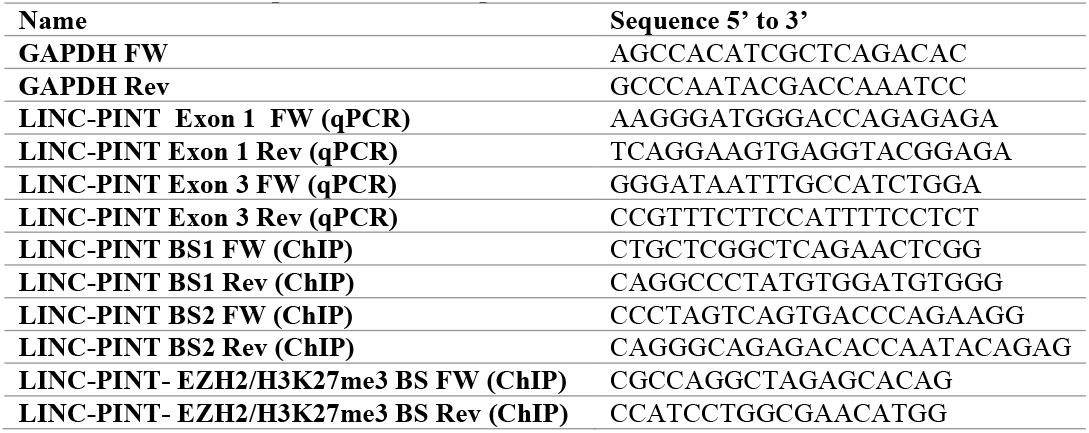
Primers for qPCR and ChIP qPCR

### Chromatin extraction

Cells were cross-linked using 1% formaldehyde (Sigma) added directly to the medium, and incubated on a shaking platform for 10 min at room temperature. The cross-linking reaction was stopped by adding 0.125 M glycine for 5 min. Cells were harvested and washed twice with PBS and then lysed in 1 ml cell lysis buffer (20 mM Tris-HCl pH 8, 85 mM KCl, 0.5% Nonidet P-40, 1:100 protease inhibitor cocktail) for 10 min on ice. Nuclear pellets were resuspended in 200 μl nuclei lysis buffer (50 mM Tris-HCl pH 8, 10 mM EDTA, 1% SDS, 1:100 protease inhibitor cocktail) for 10 min on ice, and then sonicated (Bioruptor, Diagenode) at high power settings for 3 cycles, 6 min each (30 sec ON/OFF). Samples were centrifuged (20,000g, 15 min, 4°C) and the soluble chromatin fraction was collected. In some experiments a Micrococal-Chromatin extraction protocol was used: Cells were harvested and resuspended in Buffer A (10 mM HEPES pH 7.9, 10 mM KCl, 1.5 mM MgCl_2_, 0.34 M sucrose and 10% glycerol) supplemented with 0.1% triton X-100, 1 mM DTT, 1:200 protease inhibitor mixture (PI) and 100 nM PMSF (Sigma). Cells were incubated for 8 min on ice, then centrifuged 5 min at 1850*g*, 4°C. The pellet was washed once with Buffer A supplemented with DTT, PI and PMSF, then lysed with Buffer B (3 mM EDTA and 0.2 mM EGTA) supplemented with DTT and PI, for 30 min on ice. Samples were centrifuged 5 min at 1850*g*, 4°C to pellet the chromatin fraction. Finally, chromatin fraction was solubilized in Buffer A with 1:100 micrococcal nuclease enzyme (NEB) and incubated for 15 min at 37°C shaker.

For protein-protein interaction analysis, the soluble chromatin was precleared with Magna ChIP™ Protein A+G Magnetic Beads (Millipore, 16-663) for 1h and then incubated overnight at 4°C with magnetic FLAG-M2 beads. The immunoprecipitated complexes were washed once with TSE150 buffer [20 mM tris-HCl (pH 8), 2 mM EDTA, 1% Triton X-100, 0.1% SDS, and 150 mM NaCl], TSE500 buffer [20 mM tris-HCl (pH 8), 2 mM EDTA, 1% Triton X-100, 0.1% SDS, and 500 mM NaCl], buffer 3 [250 mM LiCl, 10 mM tris-HCl (pH 8), 1 mM EDTA, 1% sodium deoxycholate, and 1% Nonidet P-40], and twice with TE buffer [10 mM tris-HCl (pH 8) and 1 mM EDTA]. Immunoprecipitated complexes were resolve in protein sample buffer and analyzed by Western blot.

### Chromatin immunoprecipitation(ChIP)-qPCR and

The chromatin fraction was diluted 5× in dilution buffer [20 mM tris-HCl (pH 8), 2 mM EDTA, 150 mM NaCl, 1.84% Triton X-100, and 0.2% SDS]. Chromatin was precleared overnight at 4°C with A+G Magnetic Beads. The precleared sample was then immunoprecipitated with magnetic FLAG-M2 beads or A/G magnetic beads preconjugated with the indicated antibody. The immunoprecipitated complexes were washed according to the chromation extraction protocol detailed above. DNA was eluted with elution buffer (50 mM NaHCO3, 140 mM NaCl, and 1% SDS) containing ribonuclease A (0.2 μg/μl) and proteinase K (0.2 μg/μl). Last, the DNA eluates were de-cross-linked at 65°C overnight with shaking at 900 rpm and purified by NucleoSpin Gel and PCR Clean-up kit (Macherey-Nagel), according to the manufacturer’s instructions. Purified DNA was subjected to qPCR using specific primers (**Table 2**). Primers for TWIST1 binding sites were designed on the basis of H3K4me3, H3K27Ac and TF clusters in LINC-PINT locus and the occurrence of E-box elements (^5^’CANNTG^3^’). Primers for EZH2 and H3K27me3 binding sites were designed using ChIP-seq data previously published (38,39) and viewed using Integrated Genomics Viewer software (40). qPCR was preformed using SYBR Green I Master (Roche) in a LightCycler 480 System (Roche). All samples were amplified in triplicate in a 384-well plate using the following cycling conditions: 5 min at 95°C, 45 cycles of amplification; 10 s at 95°C, 10 s at 60°C, and 10 s at 72°C, followed by melting curve acquisition; and 5 s at 95°C, 1 min at 65°C and monitoring up to 97°C, and lastly cooling for 30 s at 40°C. The results were normalized to input DNA and presented as % input.

### Mass spectrometry

Sample of non-radioactive methylation assay containing 2μg His-TWIST1 and 4μg GST SETD6 were incubated with 3.2 mM SAM overnight at 30°C. An additional sample without SAM served as reference. For samples preparation (Weizmann Institute of Science), Proteins were reduced with 5 mM dithiothreitol (Sigma) for 1hr at room temperature, and alkylated with 10 mM iodoacetamide (Sigma) in the dark for 45 min at room temperature. Proteins were then subjected to digestion with trypsin (Promega; Madison, WI, USA) overnight at 37°C at 50:1 protein:trypsin ratio, followed by a second trypsin digestion for 4 hr. The digestions were stopped by addition of trifluroacetic acid (1% final concentration). Following digestion, peptides were desalted using Oasis HLB, μElution format (Waters, Milford, MA, USA). The samples were vacuum dried and stored in −80°C until further analysis. ULC/MS grade solvents were used for all chromatographic steps. Each sample was loaded using split-less nano-Ultra Performance Liquid Chromatography (10 kpsi nanoAcquity; Waters, Milford, MA, USA). The mobile phase was: A) H2O + 0.1% formic acid and B) acetonitrile + 0.1% formic acid. Desalting of the samples was performed online using a reversed-phase Symmetry C18 trapping column (180 μm internal diameter, 20 mm length, 5 μm particle size; Waters). The peptides were then separated using a T3 HSS nano-column (75 μm internal diameter, 250 mm length, 1.8 μm particle size; Waters) at 0.35 μL/min. Peptides were eluted from the column into the mass spectrometer using the following gradient: 4% to 30%B in 50 min, 30% to 90%B in 5 min, maintained at 90% for 5 min and then back to initial conditions. The nanoUPLC was coupled online through a nanoESI emitter (10 μm tip; New Objective; Woburn, MA, USA) to a quadrupole orbitrap mass spectrometer (Q Exactive Plus, Thermo Scientific) using a FlexIon nanospray apparatus (Proxeon). Data was acquired in data dependent acquisition (DDA) mode, using a Top20 method. MS1 resolution was set to 70,000 (at 400 m/z), mass range of 300-1650 m/z, AGC of 3e6 and maximum injection time was set to 20 msec. MS2 resolution was set to 17,500, quadrupole isolation 1.7 m/z, AGC of 1e6, dynamic exclusion of 30 sec and maximum injection time of 60 msec. Data was analysed using Byonic search engine (Protein Metrics) against the Human protein database (SwissProt Dec20) allowing for the following modifications: fixed carbamidomethylation on C, variable protein N-terminal acetylation, oxidation on M, deamidation on NQ, methylation on K, demethylation on K and trimethylation on K. Protein FDR was set to 1%.

### RNA-seq and data processing

Total RNA was extracted from U251 cells (SETD6 control vs. KO or Empty vs. TWIST1 WT) using the NucleoSpin RNA Kit (Macherey-Nagel). Samples were prepared in triplicates (SETD6 KO) or duplicates (TWIST1-WT cells). RNA-seq libraries were prepared at the Crown Genomics institute of the Nancy and Stephen Grand Israel National Center for Personalized Medicine, Weizmann Institute of Science. Libraries were prepared using the INCPM-mRNA-seq protocol. Briefly, the polyA fraction (mRNA) was purified from 500 ng of total input RNA followed by fragmentation and the generation of double-stranded cDNA. After Agencourt Ampure XP beads cleanup (Beckman Coulter), end repair, A base addition, adapter ligation and PCR amplification steps were performed. Libraries were quantified by Qubit (Thermo fisher scientific) and TapeStation (Agilent). Sequencing was done on a Hiseq instrument (Illumina) using two lanes of an SR60_V4 kit, allocating 20M reads per sample (single read sequencing).

Data processing of SETD6 KO RNA-seq: Adaptor removal and bad quality filtering was performed using Trimmomatic-0.32. Reads were than mapped to the human genome version GRCh38 using STAR-2.3.0. Counting was done using HTSeq-count version 0.6.1. (41). Statistical analysis was done using the DESeq2 R package while normalized counts were generated using the vsd function.

Data processing for TWIST1 U251 RNA-seq: Poly-A/T stretches and Illumina adapters were trimmed from the reads using cutadapt (DOI: https://doi.org/10.14806/ej.17.1.200); resulting reads shorter than 30bp were discarded. Reads were mapped to the Homo Sapiens GRCh38 reference genome using STAR (42), supplied with gene annotations downloaded from Ensembl (and with EndToEnd option and outFilterMismatchNoverLmax was set to 0.04). Expression levels for each gene were quantified using htseq-count (41), using the gtf above. TPM values were estimated independently using Kallisto (43). Raw gene counts were normalized and compared using DESeq 2 1.23.0 (44). Differentially expressed genes (P-adj <0.05) were then subjected to hierarchical clustering using Heatmapper web tool (45).

#### Bioinformatic analysis

Kaplan-Meier survival curve and GO analysis of GBM patients was generated using the GlioVis data portal (46), based on the CGGA (47) and TCGA (https://www.cancer.gov/tcga) databases. For RNA-seq, Poly-A/T stretches and Illumina adapters were trimmed from the reads using cutadapt (DOI: https://doi.org/10.14806/ej.17.1.200); resulting reads shorter than 30bp were discarded. Reads were mapped to the Homo Sapiens GRCh38 reference genome using STAR (42), supplied with gene annotations downloaded from Ensembl (and with EndToEnd option and outFilterMismatchNoverLmax was set to 0.04). Expression levels for each gene were quantified using htseq-count (41), using the gtf above. TPM values were estimated independently using Kallisto (43). Raw gene counts were normalized and compared using DESeq 2 1.23.0 (44). Differentially expressed genes (p adj <0.05) were then subjected to hierarchical clustering using Heatmapper web tool (45). For SETD6 control vs. KO experiment, Gene set enrichment analysis (48,49) of all genes was performed for Hallmark gene sets. SETD6 and TWIST1 shared target genes were analyzed using the DAVID tool for gene ontology (GO) biological processes (50,51). For differentially expressed lncRNAs, gene symbols including “LINC” or “-AS1” were selected. Shared lncRNAs of SETD6 and TWIST1 were identified using venn diagram (http://bioinformatics.psb.ugent.be/webtools/Venn/). LINC-PINT gene region was analyzed using the UCSC genome browser for promoter region (H3K4me3), regulatory elements (H3K27Ac) and TF clusters. 210 common target genes for SETD6 and TWIST1 were analyzed in the Enrichr database for ENCODE TF ChIP-seq and Epigenomics Roadmap HM ChIP-seq. 33 common target genes of SETD6, TWIST1 and LINC-PINT were analyzed using the DAVID tool for GO biological processes.

#### Statistical analyses

Statistical analyses for all assays were performed with GraphPad Prism software, using one-way or two-way analysis of variance (ANOVA) with a Tukey’s post hoc test.

## Results

### SETD6 levels correlate with genes related to ECM organization and EMT in GBM

We utilized the GlioVis bioinformatic tool (http://gliovis.bioinfo.cnio.es/) to study the survival rate of GBM patients with high or low expression of SETD6. Kaplan-Meier survival analysis of 633 glioma patients taken from the CCGA database with high (n=313) or low (n=320) expression of SETD6 revealed that the expression of this methyltransferase can serve as a predictor for overall survival in these patients. Specifically, patients with high expression of SETD6 had a significantly lower (p=0.0011) survival rate compared to patients with low expression of SETD6 (**Figure 1A**). Complementary to these findings, Gene Ontology (GO) analysis of TCGA data (https://www.cancer.gov/tcga.) further suggests that SETD6 expression level in GBM patients associated with genes that regulate ECM organization (**Figure 1B**). These findings imply that SETD6 may have clinical significance in the pathobiology of GBM. To study the potential role of SETD6 in GBM, we next performed an RNA-seq experiment using U251 control (CT) and two SETD6 CRISPR knock-out (KO) cells derived from two independent gRNAs clones (**Figure 1C**). Sequence validation of the two KO cells is shown in **supplementary figure 1A**. In this analysis, 2104 differentially expressed genes were identified (p adj <0.05); of these, 1190 genes were down-regulated and 914 were up-regulated in the CRISPR SETD6 KO cells (**Figure 1C**). Gene set enrichment analysis (GSEA) of hallmark gene sets revealed a significant enrichment of EMT in the down-regulated genes which is known to be a key regulatory process in GBM (52-54) (**Figure 1D and 1E**). For the up-regulated genes, we observed less significant results, with an enrichment of pathways linked to inflammation and oncogenic processes (**Supplementary figure 1B**), processes in which we and others have shown the involvement of SETD6 (25,55). ECM organization is tightly linked to EMT. Of relevance, EMT is known to contribute to the aggressive phenotype of GBM (56-59). We then hypothesize that SETD6 may regulate GBM progression by a direct or indirect effect on EMT related processes.

**Figure1.**
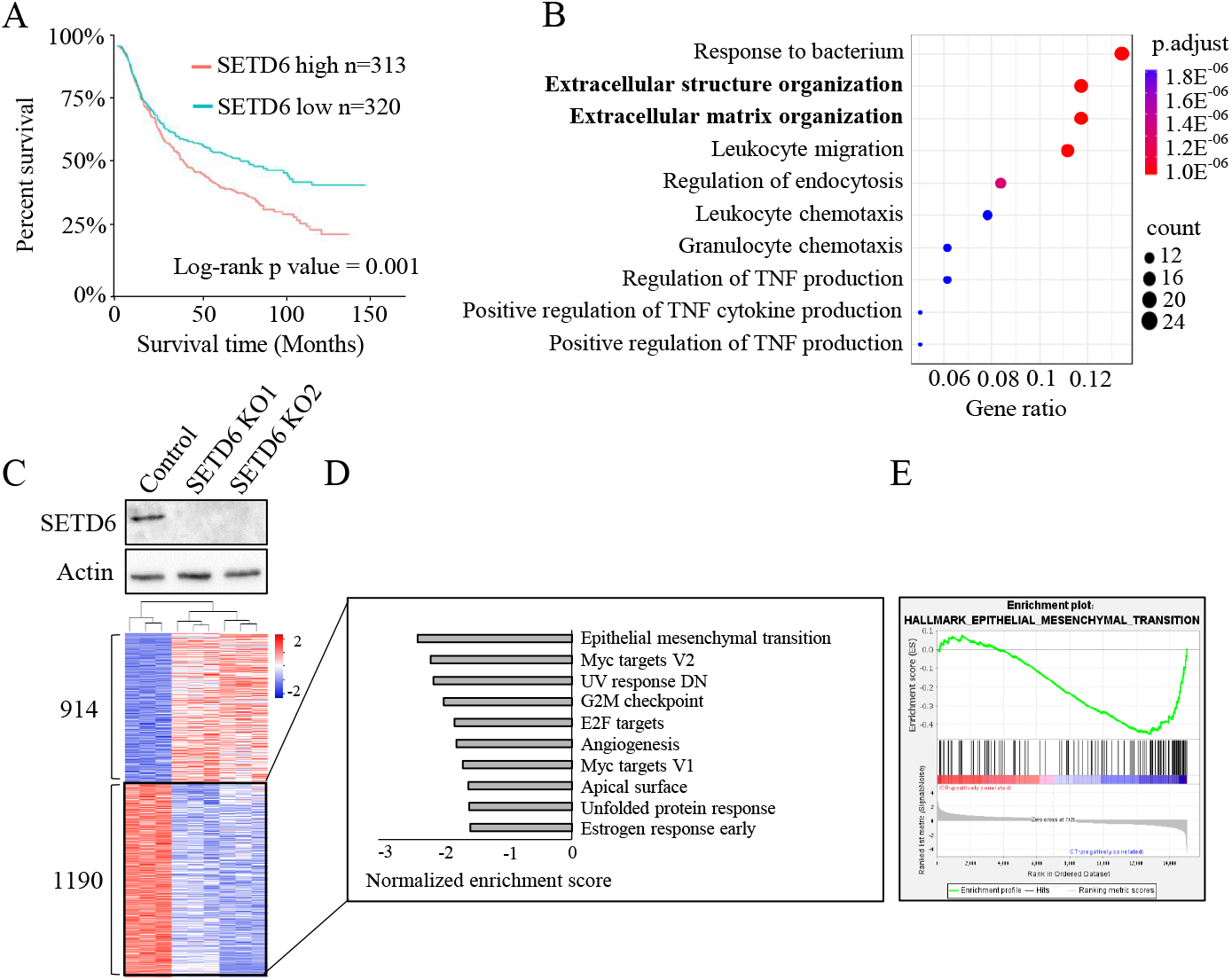
SETD6 levels correlates with genes related to ECM organization and EMT in GBM. (A) Kaplan-Meier survival curve of GBM patients stratified by high and low SETD6 expression levels. The data was generated using the CGGA database and the GlioVis data portal (GlioVis data portal for visualization and analysis of brain tumor expression datasets. (B) Gene ontology (GO) analysis showing biological processes related to SETD6 in GBM. The data was generated using the TCGA database and the GlioVis data portal. (C) U251 control and CRISPR SETD6 KO cells (two independent gRNAs) were subjected to western blot analysis using SETD6 and Actin antibodies (Top). Bottom: Heatmap showing up- and down-regulated genes (p adj < 0.05) from RNA-seq analysis of the indicated cells. Red and blue colors represent high and low expression levels, respectively. (D) Differentially expressed genes were analyzed using the Gene set enrichment analysis (GSEA) platform. Hallmark gene sets enriched in genes down-regulated in SETD6 KO cells are presented according to their normalized enrichment score. (E) Enrichment plot of EMT gene set. The y-axis shows the enrichment score for each gene in the gene set (vertical black line represents each gene). Red (high) and blue (low) represent the expression levels in SETD6 KO cells vs. control cells.

### SETD6 binds and methylates TWIST1 in-vitro and in cells

The transcription factors SLUG, SNAIL, TWIST1 and ZEB1 are major regulators of the EMT process in GBM (10,60). We hypothesized that SETD6 may have functional link to one or more of these factors. To this end, we first expressed and purified the recombinant proteins of SLUG, SNAIL and TWIST1. For technical reasons, we were unable to purify ZEB1 (data not shown). We first assessed the potential direct interactions between SETD6 and these proteins using ELISA. A direct significant interaction was observed between SETD6 and TWIST1, no interaction was observed between SETD6 and SNAIL, and a moderate but significant interaction was seen with SLUG. GST and BSA served as negative controls for these experiments (**Figure 2A**). To further analyze the interaction of SETD6 and TWIST1 in cells, we co-transfected HA-SETD6 and FLAG-TWIST1 in HEK293T cells followed by immunoprecipitation. As shown in **Figure 2B**, SETD6 interacts with immunoprecipitated TWIST1. Since TWIST1 is a transcription factor and localized primarily to the nucleus (61) and due to the established role of SETD6 in transcription regulation (26,28,31) we hypothesized that the interaction takes place on chromatin. Co-immunoprecipitation experiments within an isolated chromatin fraction in SETD6-KO HEK293T cells confirmed that the two proteins interact at chromatin (**Figure 2C**).

**Figure 2.**
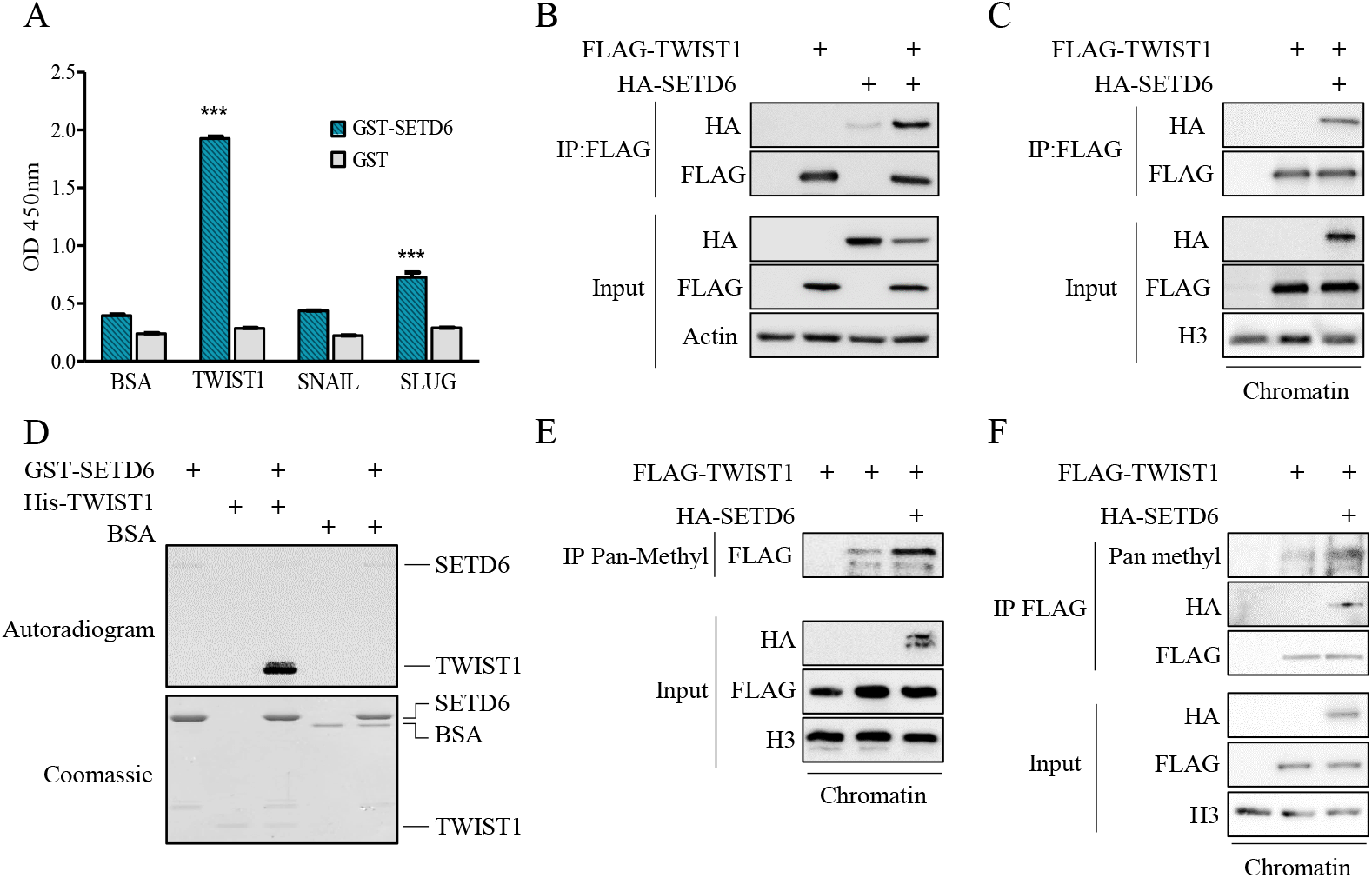
SETD6 interacts and methylates TWIST1 *in-vitro* and in cells. (A) Enzyme-linked immunosorbent assay (ELISA) was performed with the indicated recombinant proteins. The graph represents absorbance at 450nm for each condition. (B+C) Human embryonic kidney (HEK) 293T cells were transfected with FLAG-TWIST1 with or without HA-SETD6. Whole cell lysates (B) or chromatin fraction (C) were immunoprecipitated with FLAG-M2 beads, followed by Western blot analysis with indicated antibodies. (D) *In-vitro* methylation of TWIST1 by SETD6. Samples were subjected to SDS–polyacrylamide gel electrophoresis (PAGE) followed by exposure to autoradiogram to detect 3H-labeled proteins or Coomassie staining to detect all proteins. BSA used as a negative control. (E+F) U251 cells were transfected with FLAG-TWIST1 with or without HA-SETD6. Chromatin fractions were immunoprecipitated with Pan-Methyl lysine antibody (E) or FLAG-M2 beads (F), followed by Western blot analysis with the indicated antibodies.

Given the enzymatic activity of SETD6 (62,63) and its physical interaction with TWIST1 *in-vitro* and in cells, we hypothesized that SETD6 methylates TWIST1. In an *in-vitro* methylation assay containing recombinant His-TWIST1, GST-SETD6 and tritium labeled SAM (S-adenosyl-methionine, the methyl donor), we found that SETD6 methylates TWIST1 and not the negative control BSA (**Figure 2D**). We could not detect any methylation of SNAIL. Furthermore, a weak methylation signal was detected for SLUG (**Supplementary figure 2**) which is consistent with the results shown in Figure 2A. To validate whether SETD6 methylates TWIST1 in cells, we extracted the chromatin fraction of U251 cells overexpressing exogenous FLAG-TWIST, with or without exogenous HA-SETD6 overexpression, followed by immunoprecipitation with pan-methyl antibody. We found that the methylation of TWIST1 at chromatin in U251 cells was increased in the presence of SETD6 (**Figure 2E**). The weak signal in the absence of SETD6 (lane 2), suggests that TWIST1 is methylated by endogenous SETD6 or by an additional methyltransferase. To further validate that SETD6 methylates TWIST1 at chromatin, we performed a reciprocal experiment in which FLAG-TWIST1 was immunoprecipitated from U251 chromatin extracts. As shown in **Figure 2F**, TWIST1 methylation increased in the presence of SETD6 overexpression. Taken together, our results show that SETD6 binds and methylates TWIST1 *in-vitro* and in cells at the chromatin.

### TWIST1 function in cell adhesion and migration is SETD6 dependent

Given the transcriptional activity mediated by SETD6 in U251 shown in Figure 1C and the methylation of TWIST1 by SETD6, we hypothesized that both proteins participate in the regulation of similar transcriptional programs. To address this, we have performed RNA-seq experiment in U251 cells stably expressing Flag-TWIST1 (**Figure 3A**) and then crossed the data with the differentially expressed genes regulated by SETD6. A total of 1075 genes were differentially expressed (p < 0.05) between FLAG-TWIST1 to empty vector expressing cells. As expected, EMT and extracellular matrix related-processes were highly enriched in FLAG-TWIST1 expressing cells (**Supplementary Figure 3**). Among the 1075 genes, 210 SETD6 and TWIST1 shared genes were identified (P-value <1.6E^-19^) (**Figure 3B**). In a gene ontology (GO) analysis, we identified significant enrichment of processes related to EMT such as ECM organization, collagen organization, cell adhesion and migration (**Figure 3C**). Interestingly, these biological processes are correlated with the GO terms extracted from the mRNA data of GBM patients presented in Figure 1. We therefore asked if TWIST1 expression regulates these processes in a SETD6-dependent manner. In a cell adhesion assay performed on fibronectin (an ECM protein) coated wells, we observed loss of cell adhesion in cells stably expressing TWIST1. However, when TWIST1 was expressed in SETD6-depleted cells, we observed a similar level of cell adhesion as the control cells (**Figure 3D and 3E**). Loss of cell adhesion is tightly linked to increased migration (64). Consistent with that, in wound healing (**Figure 3F and 3G**) and transwell migration assays (**Figure 3H**), SETD6 knockout significantly reduced migration abilities of cells stably expressing TWIST1 compared to those expressing TWIST1 without SETD6 knockout. Together, our results suggest that TWIST1 induction of loss of cell adhesion and increased migration are SETD6 dependent.

**Figure 3.**
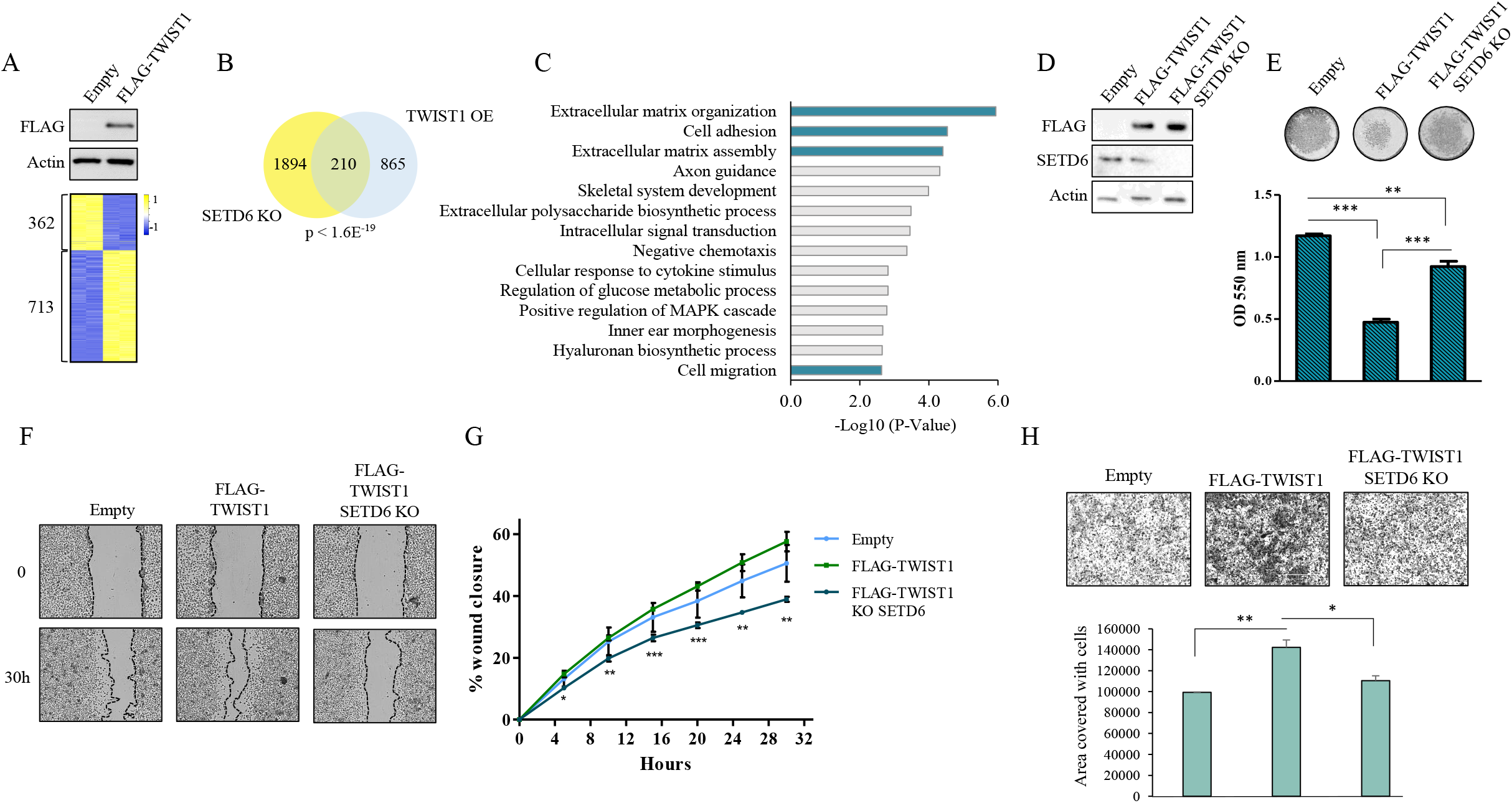
TWIST1 function in cell adhesion and migration is SETD6 dependent. (A) U251 stably expressing empty vector or FLAG-TWIST1 were subjected to Western blot analysis with the indicated antibodies (Top). Heatmap showing up-and down-regulated genes (p-value <0.05) from RNA-seq analysis of the indicated cells. color bar represents high (yellow) and low (blue) expression levels. (B) Venn diagram showing common genes for TWIST1 and SETD6 as identified in the RNA-seq analyses. (C) Common genes were analyzed using the DAVID database. The most significantly enriched biological processes are presented (p-value<0.05) and relevant processes are highlighted in cyan. (D) U251 control or SETD6 KO cells stably expressing empty vector or FLAG-TWIST1 were subjected to Western blot analysis with the indicated antibodies. (E) Fibronectin adhesion assay with the indicated cells. Top: representative images of fibronectin-adherent cells stained with Crystal violet. Bottom: Crystal violet stained cells were dissolved in 2% SDS and the absorbance at 550nm was measured. (F+G) Wound healing assay with the indicated cells. Confluent cells were scratched with 200ul pipette tip. Wound closure was monitored and calculated by Lionheart™ FX automated microscope and representative images at 0 and 30h are shown with black lines indicating wound edges (Left). Right: % wound closure (mean+SEM) of each cell type is shown. Statistical significance of each time point was calculated using two-way ANOVA (*p<0.05, ** p<0.01, *** p<0.001). (H) Transwell migration assay. Serum starved cells were seeded onto a semi-permeable membrane inserts with 10% serum medium below. After 24h, migrated cells were fixed and stained and images were taken using EVOS FL Cell Imaging System and analyzed using Fiji software. Representative images are shown (Top). The graph (Bottom) represents mean area covered with migrated cells of 4 fields for each cell type. Statistical significance was calculated using one-way ANOVA (*p<0.05, ** p<0.01).

### SETD6 methylates TWIST1 on lysine 33

In order to map the methylation site, we performed non-radioactive methylation assay followed by mass spectrometry analysis. Among the 10 lysine residues found in TWIST1, lysine-33 and lysine-73 were identified as methylated by the mass spectrometry analysis (**Figure 4A and 4B**), however the K73 methylation was also detected in the sample with no SAM (negative control). For validation, we have generated methylation-deficient TWIST1 mutants at lysine-33 and lysine-73 to arginine (K33R, K73R) using site-directed mutagenesis. In a radioactive methylation assay in the presence of GST-SETD6 with either WT TWIST1 or K33R, K73R mutants, we observed a decrease in the methylation signal for K33R mutant but not for K73R mutant (**Figure 4C**). In addition, we found that immunoprecipitated K33R TWIST1 from chromatin fraction of HEK293T cells is less methylated *in-vitro* by recombinant SETD6 compared to WT TWIST1 (**Figure 4D**). We next generated a site-specific antibody for TWIST1 K33me which specifically recognized a TWIST1 mono-methylated peptide at K33 but not the unmodified one (**Figure 4E**). Moreover, using this antibody, we could confirm the methylation of stable over-expressed TWIST1 WT in U251 but not TWIST1 harboring a K33R mutation (**Figure 4F**). Taken together, these results suggest that lysine 33 is the primary methylation site.

**Figure 4.**
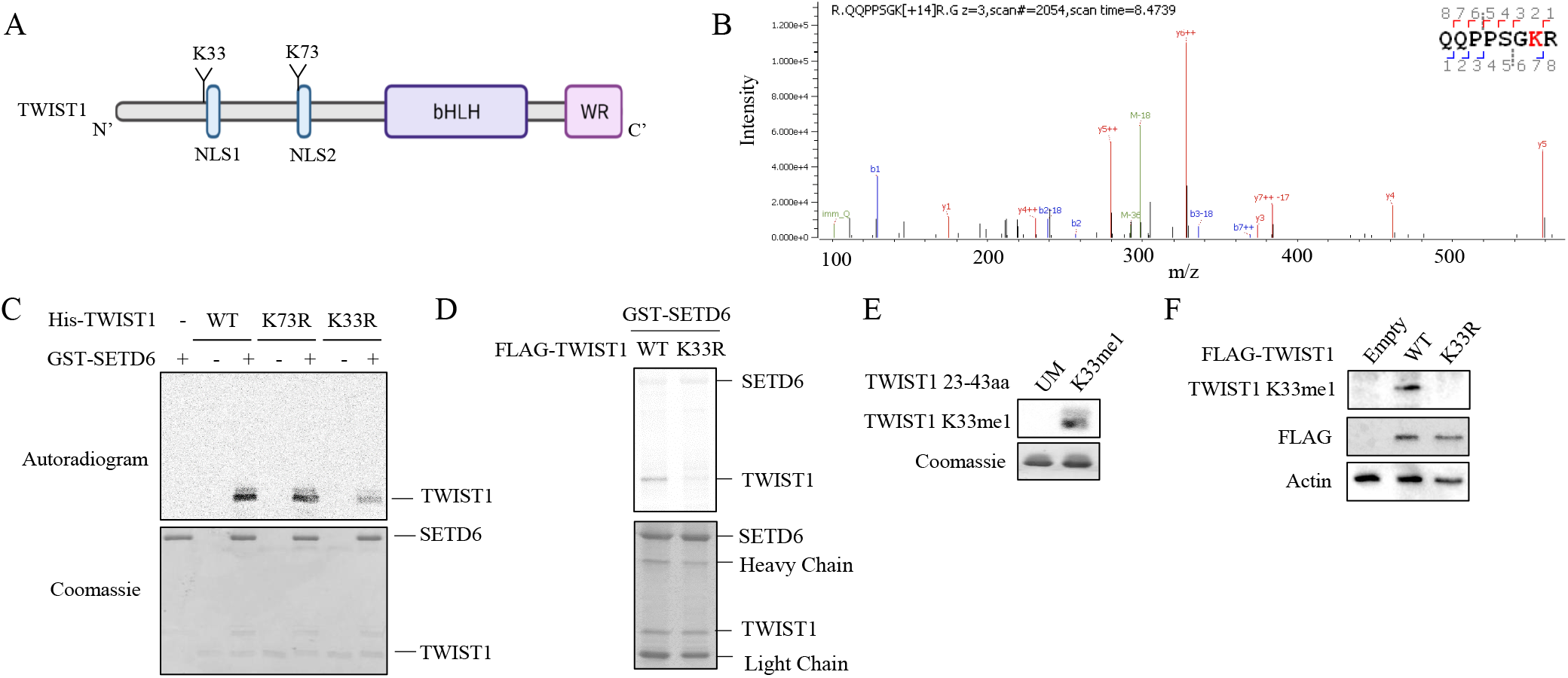
SETD6 methylates TWIST1 at K33. (A) Illustration of TWIST1 sequence and domains with the identified methylated lysine residues. (B) MS spectra of TWIST1 QQPPSGKR peptide. In-vitro methylation assay was performed with recombinant His-TWIST1 and GST-SETD6 with or without SAM followed by mass spectrometry analysis. (C) MS results were validated by *in-vitro* methylation assay with the indicated recombinant proteins in the presence of ^3^H-labeled SAM. (D) semi-*in-vitro* methylation assay. HEK293T SETD6 KO cells were transfected with FLAG-TWIST1 WT or K33R mutant. Chromatin fractions were immunoprecipitated with FLAG-M2 beads and were subjected to radioactive *in-vitro* methylation assay with recombinant SETD6. Autoradiogram used to detect ^3^H-labeled proteins and Coomassie staining to detect all proteins. (E) TWIST1 peptides (Unmodified and K33me1) were subjected to SDS-PAGE followed by western blot analysis with K33me1 antibody and coomassie staining. (F) Western blot analysis of U251 stably expressing empty vector, FLAG-TWIST1 WT or K33R cells with the indicated antibodies.

### Methylated TWIST1 is enriched at the long non-coding RNA LINC-PINT locus

Our RNA-seq data revealed that both SETD6 and TWIST1 regulate the expression of several long non-coding RNAs (Lnc-RNAs) (**Figure 5A**). Among the 28 common Lnc-RNAs, 8 had a fold change of more than 1.5 (**Figure 5B**). We hypothesized that SETD6-dependent TWIST1 methylation might regulate the expression of these lnc-RNAs. We have decided to focus on LINC-PINT since previous studies connected its expression with inhibition of cell migration and invasion in several cancer types, including GBM (34,35,65,66).

**Figure 5.**
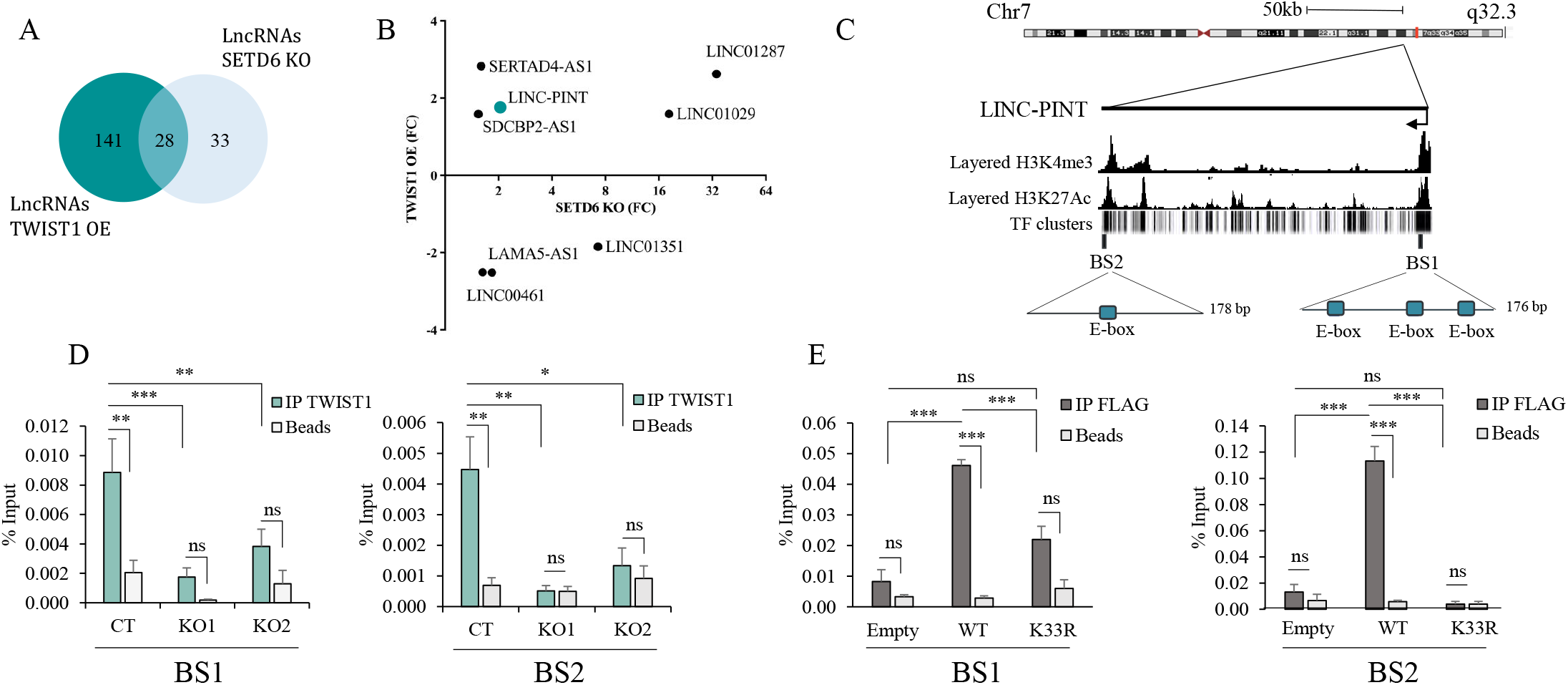
Methylated TWIST1 binds LINC-PINT gene region. (A) Venn diagram showing lncRNAs regulated by both SETD6 and TWIST1, which identified in the previous RNA-seq experiments. (B) Graph of top 8 shared lncRNAs (FC>1.5) showing their FC in SETD6 KO cells (x-axis) and cells stably express FLAG-TWIST1 (y-axis). (C) LINC-PINT genomic region. H3K4me3, H3K27Ac and TF clusters were extracted from the UCSC genome browser. The two TWIST1 binding sites (BS1 and BS2) used for ChIP assays are shown. E-boxes inside each binding site are indicated. (D) ChIP assays of U251 SETD6 KO cells immunoprecipitated with TWIST1 antibody or beads as negative control. (E) ChIP assay of U251 stably expressing empty vector, FLAG-TWIST1 WT or K33R cells immunopercipitated with FLAG-M2 beads. (D+E) Graphs show % input of the quantified DNA. Error bars are SEM. Statistical significance was calculated using two-way ANOVA for 3 experimental repeats (ns, non-significant, * p<0.05, **p<0.01, ***p<0.001).

By exploring the genomic location of LINC-PINT on chromosome 7q32.3, we have identified 2 potential TWIST1 binding sites (BS1 and BS2) by searching for E-box elements (**Figure 5C**). Using chromatin immunoprecipitation (ChIP) experiments, we revealed enrichment of endogenous TWIST1 on BS1 and BS2, confirming the hypothesis that TWIST1 binds the LINC-PINT locus in these locations (**Figure 5D**). In contrast, the occupancy of endogenous TWIST1 was significantly reduced in SETD6 KO cells at both binding sites, suggesting that TWIST1 occupancy is SETD6 dependent (**Figure 5D**). Consistent with these findings, the occupancy of FLAG-TWIST1 WT was significantly higher compared to Flag-TWIST1 K33R mutant at BS1 and BS2 regions (**Figure 5E**), indicating that the presence of TWIST1 on LINC-PINT locus is dependent on K33 methylation.

### TWIST1 methylation negatively regulates the expression of LINC-PINT

Based on the observations that LINC-PINT expression is augmented and TWIST occupancy at the LINC-PINT locus is reduced under conditions of either TWIST-K33R expression or SETD6 KO, our working hypothesis was that this methylation event regulates the expression of LINC-PINT. To address this hypothesis, we first tested by qPCR the expression of LINC-PINT and found a significant increase under conditions of SETD6 KO compare to control cells (**Figure 6A**). Consistent with these results, an increase in LINC-PINT levels was seen in cells stably expressing TWIST1 K33R mutant compared to TWIST1 WT, while the expression of LINC-PINT was significantly reduced in cells expressing TWIST1 WT (**Figure 6B**). Together, our results suggest that methylated TWIST1 binds the LINC-PINT locus and inhibits its expression in a SETD6 and a methylation dependent manner.

**Figure 6.**
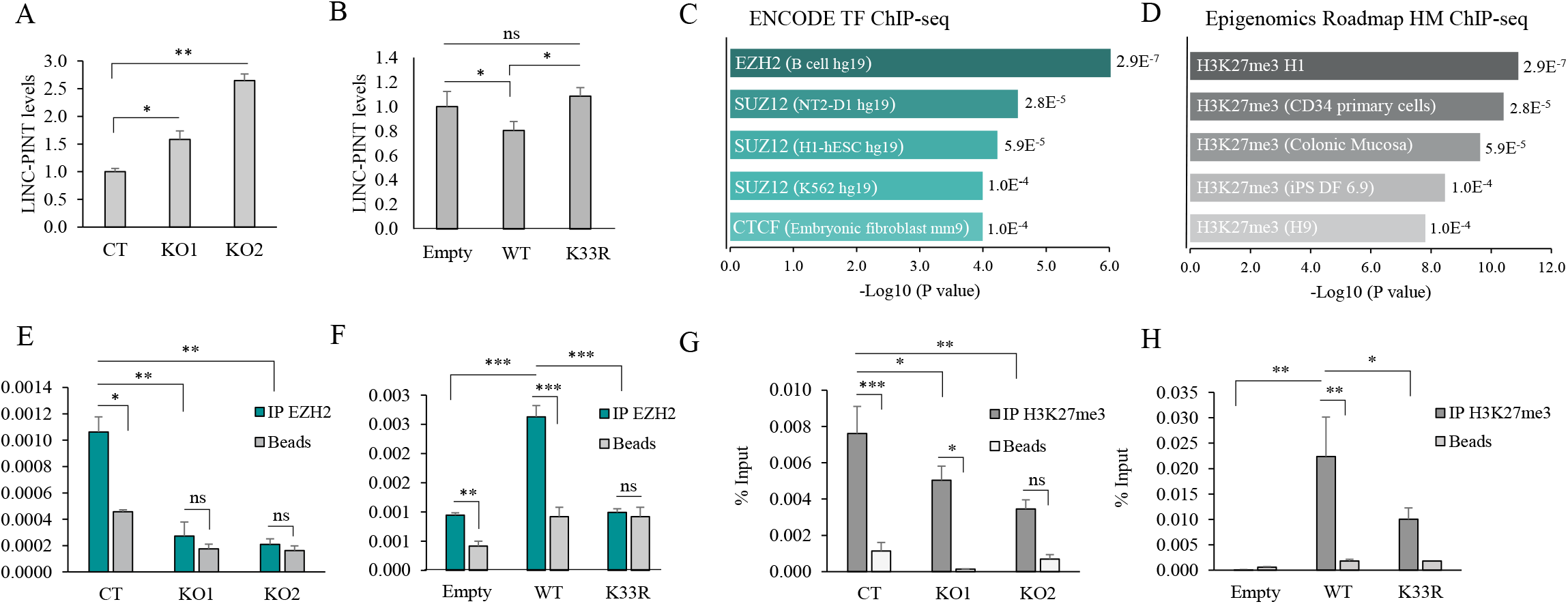
Methylated TWIST1 inhibit LINC-PINT expression. (A) RNA was extracted from U251 SETD6 KO or U251 stably expressing empty vector, FLAG-TWIST1 WT or K33R cells (B) and expression level were measured by quantitative polymerase chain reaction (qPCR). mRNA levels were normalized to glyceraldehyde-3-phosphate dehydrogenase (GAPDH) and then to empty cells. Error bars are SEM. Statistical analysis was performed for three experimental repeats using one-way ANOVA (*P < 0.05, **P < 0.01). (C+D) Common target genes of TWIST1 and SETD6 were analyzed by the Enrichr platform. Top 5 results for ENCODE TF ChIP-seq (C) and Epigenomics Roadmap HM ChIP-seq (D) are presented. (E-H) ChIP assays. Chromatin fractions of the indicated cell type were immunopercipitated with EZH2 (E+F) or H3K27me3 (G+H) antibodies or beads as negative control, followed by qPCR of LINC-PINT locus. Graphs show % input of the quantified DNA. Error bars are SEM. Statistical significance was calculated using two-way ANOVA for 3 experimental repeats (ns, non-significant, * p<0.05, **p<0.01, ***p<0.001).

We next sought for the mechanism by which TWIST1 methylation inhibits LINC-PINT expression. To address this, we analyzed TWIST1 and SETD6 common target genes using the Enrichr platform (67-69) for ENCODE TF (http://genome.ucsc.edu/ENCODE/downloads.html) (**Figure 6C**) and Epigenomics Roadmap (http://www.roadmapepigenomics.org/data) (**Figure 6D**) ChIP-seq databases. Both platforms integrate a large collection of ChIP-seq data to predict protein interaction with the DNA (39,70). The results demonstrate a significant enrichment of the PRC2 (Polycomb repressive complex 2) components EZH2 and SUZ12 (**Figure 6C**) and a significant enrichment for H3K27me3 (**Figure 6D**), a chromatin repressive mark catalyzed by EZH2 (71), associated with gene silencing (72). A snapshot of ChIP-seq data showing the enrichment of EZH2 and H3K27me3 in a brain tissue at this genomic location is shown in **Supplementary Figure 4**. To validate these results, we performed ChIP experiments to test the occupancy of EZH2 and H3K27me3 on the LINC-PINT locus. As shown in **Figure 6E**, the enrichment of the methyltransferase EZH2 decreased in the SETD6 KO cells at the LINC-PINT locus. As predicted and consistent with our working model, EZH2 displayed significantly higher enrichment in cell expressing TWIST1 WT compared to TWIST1 K33R mutant (**Figure 6F**). Likewise, H3K27me3 was significantly lower in the SETD6 KO and in cells stably expressing TWIST1 K33R, two conditions which represent un-methylated status of TWIST1 at K33 (**Figure 6G and 6H**, respectively). Taken together, these results suggest that TWIST1 methylation at K33 by SETD6 represses LINC-PINT transcription by increasing the occupancy of EZH2 and the catalysis of H3K27me3 repressive mark.

### TWIST1 mediated-inhibition of LINC-PINT leads to loss of cell adhesion and increased migration

Integration of our RNA-seq data of SETD6 (Figure 1C) and TWIST1 (Figure 3A) target genes with previously published LINC-PINT target genes (33), revealed 33 common target genes (Figure 7A). GO analysis for biological processes found enrichment of extracellular matrix organization (**Figure 7A and 7B**). LINC-PINT was shown before to regulate cell adhesion genes and to inhibit migration of cancer cells (34,73). We therefore hypothesized that cells expressing un-methylated TWIST1 (K33R mutant), which induces the expression of LINC-PINT RNA, will display similar phenotypes to cells expressing LINC-PINT. To address this hypothesis, we have generated cells stably expressing TWIST1 WT, K33R mutant and LINC-PINT and tested adhesion and migration abilities (**Figure 7C, and Supplementary Figure 5 for expression validation**). Consistent with the results obtained in Figure 3E, we found reduced adhesion in cells expressing TWIST1 WT. However, a significant increase in cell adhesion was observed in cells expressing TWIST1 K33R and LINC-PINT (**Figure 7C**). A significant inhibition in the ability of cells to close the wound in a scratch assay was observed in cells stably expressing TWIST1 K33R and LINC-PINT compared to TWIST1 WT (**Figure 7D**). In summary, our findings demonstrate that SETD6 selectively regulates the expression of LINC-PINT RNA. SETD6-mediated methylation of TWIST1 at K33 represses the expression of LINC-PINT by increasing H3K27me3 repressive mark at the LINC-PINT locus. Under permissive conditions, when TWIST1 is not methylated (SETD6-depletion or expression of TWIST1 K33R mutant), TWIST1 dissociates from the LINC PINT locus, H3K27me3 mark is decreased allowing the increase in LINC-PINT expression level to increase cell adhesion and to reduce cell migration (**Figure 7E**).

**Figure 7.**
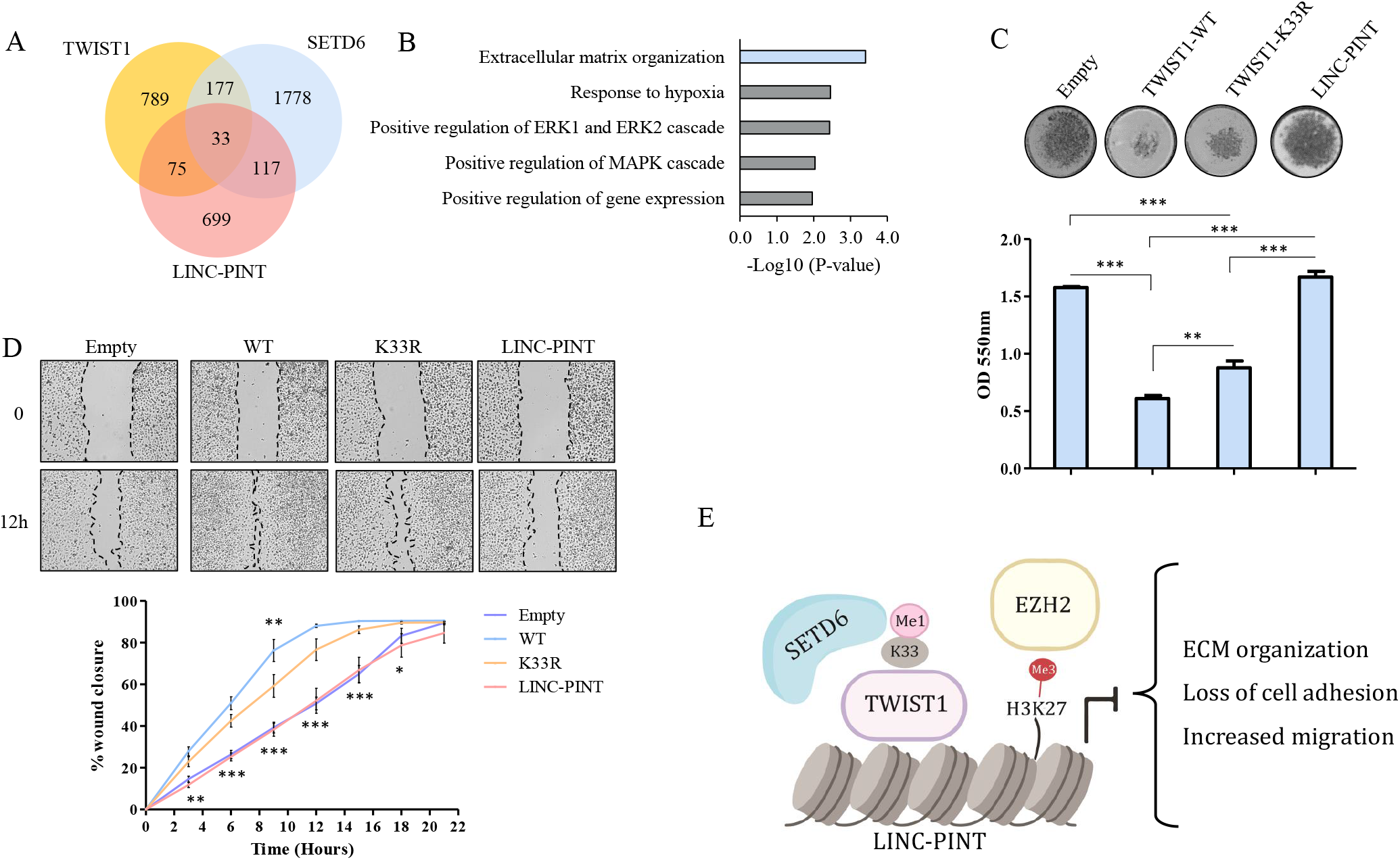
TWIST1 mediated-inhibition of LINC-PINT leads to loss of cell adhesion and increased migration. (A) Venn diagram showing common target genes for SETD6, TWIST1 and LINC-PINT. LINC-PINT target genes were taken from LINC-PINT KD HCT116 cells. (B) 33 common genes were analyzed using the DAVID database. Top 5 Significantly enriched biological processes are presented. Biological process of interest is highlighted in light blue. (C) Fibronectin adhesion assay with the indicated cells. Top: representative images of fibronectin-adherent cells stained with Crystal violet. Bottom: Crystal violet stained cells were dissolved and the absorbance at 550nm was measured. (D) Wound healing assay with the indicated cells. Confluent cells were scratched with 200ul pipette tip. Wound closure was monitored and calculated by Lionheart™ FX automated microscope and representative images at 0 and 30h are shown with black lines indicating wound edges (Top). Bottom: % wound closure (Mean+ SEM) of each time point and cell type is shown. Statistical significance was calculated using two-way ANOVA for cells stably expressing TWIST1 WT vs. LINC-PINT (below the graph) and for TWIST1 WT vs. K33R (above the graph) (*p<0.05, ** p<0.01, *** p<0.001). (E) Schematic model illustrating the inhibition of LINC-PINT expression by TWIST1 methylation. Following TWIST1 K33 methylation by SETD6, EZH2 is recruited to LINC-PINT locus and inhibits its expression through the induction of H3K27 tri-methylation. LINC-PINT inhibition leads to ECM re-organization, loss of cell adhesion and increased migration.

## Discussion

Bioinformatic analysis for SETD6 expression in GBM patients and U251 - GBM-derived cells revealed two interesting observations; First, lower expression of SETD6 correlates with better prognosis compared to patients with high SETD6 levels. Second: GO analysis for SETD6 expression from these patients and from our RNA-seq experiments using SETD6-depleted cells have suggested an enrichment of processes linked to EMT such as extracellular structure organization and cell adhesion. The findings presented in this manuscript stem from these two observations as we have hypothesized that SETD6 may regulate GBM through a functional crosstalk with one of the key cellular EMT-related transcription factors: SNAIL, SLUG, TWIST1 and ZEB. While all these transcription factors were shown to play key role in GBM (7-11), TWIST1 showed the most compelling results with regards to SETD6.

In recent years the biology of LINC-PINT has been studied in several cancer models including glioblastoma and was shown to be activated by p53 in some of them (33,34). Thus, it is not surprising that similarly to p53, its expression is downregulated in various cancers and exhibits tumor suppressor cellular properties like inhibition of proliferation, migration and invasion (34,65,66,73). Beside the several p53 binding sites along the LINC-PINT genomic locus that were characterized by others (33), here we propose that LINC-PINT expression is also transcriptionally modulated by TWIST1, and its expression is selectively regulated by the methylation status of TWIST1. This selective activation, mediated by SETD6, allows fine tuning of LINC - PINT expression and biological functions.

How the methylation of TWIST1 directly regulates the expression of LINC-PINT remains an open question. Here we provide molecular evidence that TWIST1 methylation by SETD6 at K33 increases the occupancy of EZH2 and the H3K27me3 repressive mark at the LINC-PINT locus. Similar to previous observations that LINC-PINT repression of downstream target genes is mediated by H3K27me3 (33,34), our data suggest that its own transcription regulation might be controlled by the same protein complex. An intriguing possibility, that potentially has to be validated at some point in time is that LINC-PINT regulates its own expression in a positive feedback loop mechanism. While this closed chromatin state can partially explain why LINC-PINT expression is repressed in the presence of SETD6, future biochemical and structural characterizations are required to precisely understand how TWIST1 methylation recruits these factors; what is the kinetics of this phenomenon; what are the protein complexes recruited to the LINC-PINT locus and why the lack of methylation of TWIST1 at K33 enables an elevated expression of LINC-PINT?

As described in detail in the introduction, TWIST1 is subjected to numerous post-translational modifications (18-23). However, to the best of our knowledge this is the first report which shows that it is subjected to lysine methylation. Our data suggests that TWIST1 might be also methylated on additional lysine residues in a SETD6-dependent and -independent manner. Future experiments will determine the contribution of additional methylation sites to TWIST1 cellular activity which are probably mediated by other methyl-transferases. The cross-talk between these modifications will shed new light on its cellular activities.

The correlation between SETD6 expression and the clinical outcome led us to investigate the role of this methyltransferase in GBM and enabled us to decipher a new avenue by which lysine methylation signaling controls TWIST1 activity at chromatin. Besides GBM, SETD6, TWIST1 and LINC-PINT are also expressed in several other cancers (74-76). We therefore speculate that the mechanism described in this paper can be expended and studied in other cancer models with the aim to translate these new findings for prognostic and therapeutic applications.

## Acknowledgments

We thank the Levy lab for technical assistance and helpful discussions. We thank Idan Cohen and Debbie Toiber for providing the EZH2 and H3K27me3 antibodies. We would also like to thank David Morgenstern for his help with the mass spectrometry data analysis.

## Author Contribution

LE, MF and DL conceived and designed most of the experiments. AC and KB performed the mass spectrometry analysis. TE performed the experiments to validate the TWIST1K33me1 antibody. CJF generated the TWIST1 K33me antibody. GS and NS performed the bioinformatic analysis. LE and DL wrote the paper. All authors read and approved the final manuscript.

## Supplementary Figures

**Figure S1.**
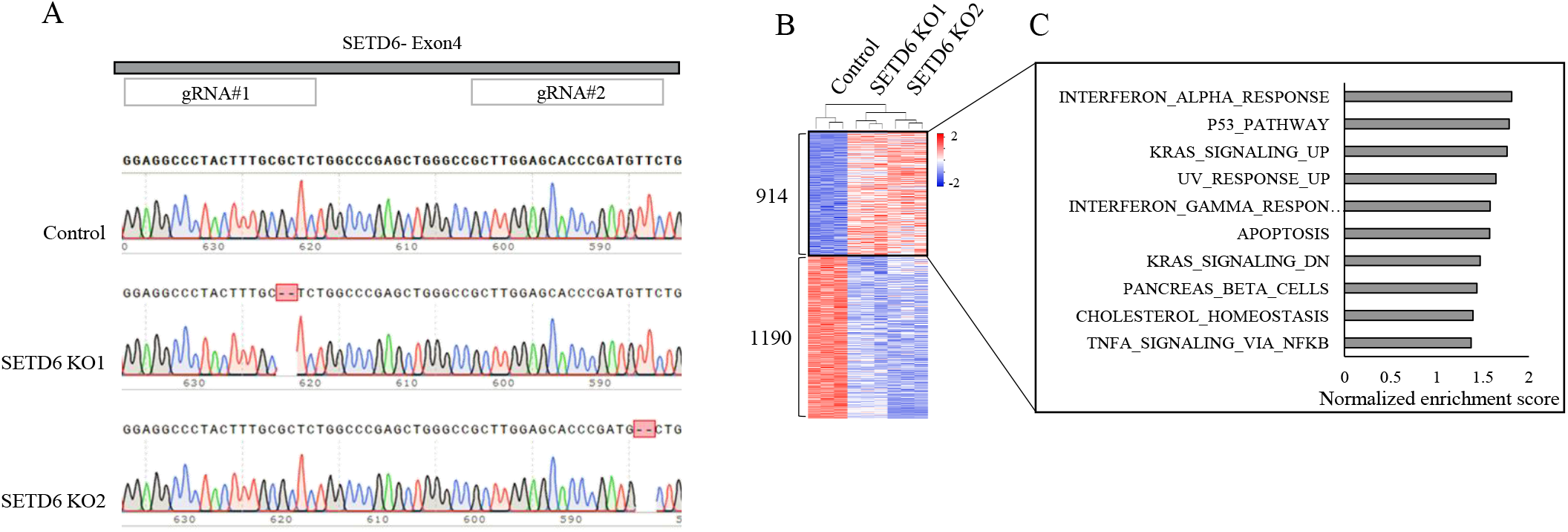
(A) Validation of CRISPR SETD6 Knock-out cells. Chromatograms of Sanger sequencing of control cells and two SETD6 knock-out cells generated from two independent gRNAs targeted for SETD6 exon 4. (B) Heatmap showing up- and down-regulated genes (p value< 0.05) from RNA-seq analysis of the indicated cells. Red and blue colors represent high and low expression levels, respectively. (C) Differentially expressed genes were analyzed using the Gene set enrichment analysis (GSEA) platform. Hallmark gene sets enriched in genes up-regulated in SETD6 KO cells are presented according to their normalized enrichment score.

**Figure S2.**
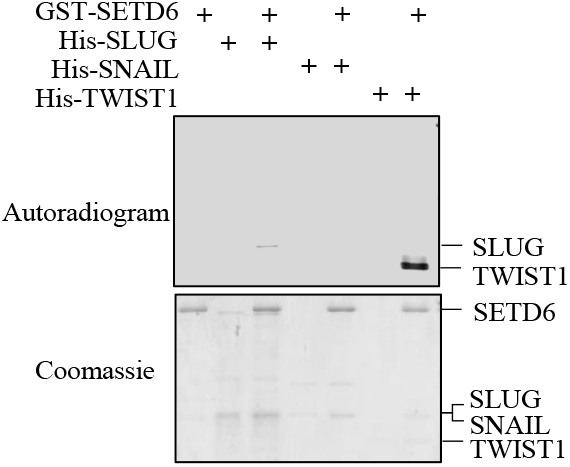
*In-vitro* methylation assay with the indicated recombinant proteins. Samples were subjected to SDS-polyacrylamide gel electrophoresis (PAGE) followed by exposure to autoradiogram to detect 3H-labeled proteins or Coomassie staining to detect all proteins.

**Figure S3.**
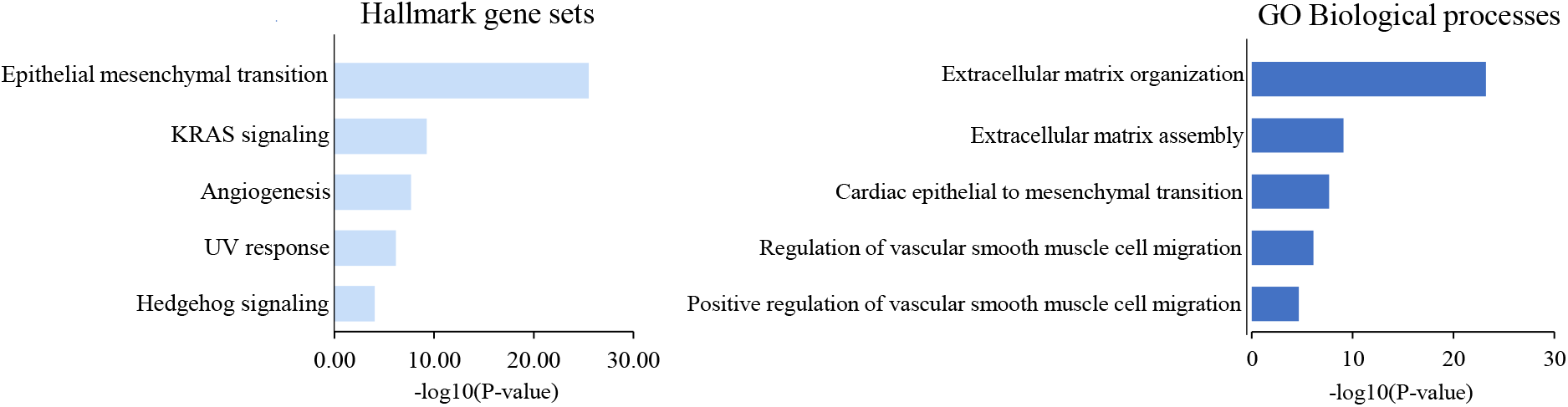
Differentially expressed genes between FLAG-WT and empty vector expressing cells were analyzed in the Enrichr platform for Hallmark gene set and GO biological processes,

**Figure S4.**
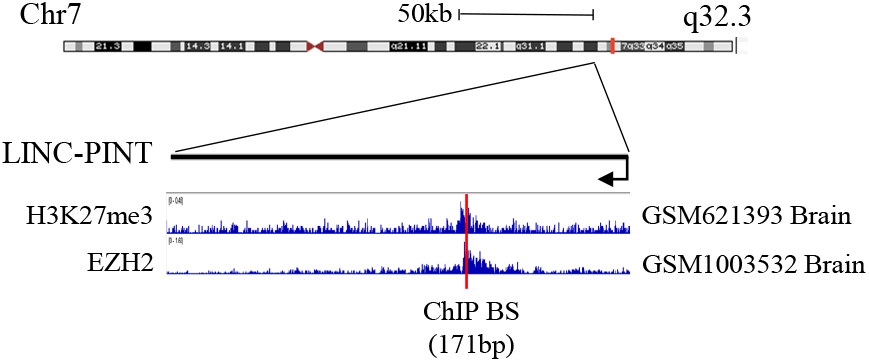
LINC-PINT genomic region with EZH2 and H3K27me3 ChIP-seq data and the region used for ChIP assays presented as red line. Data were extracted using the Cistrome data browser (http://cistrome.org/db) and visualized using the IGV software.

**Figure S5.**
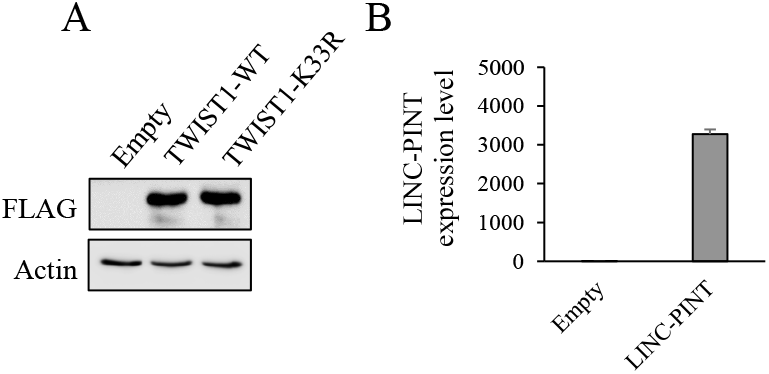
(A) U251 stably expressing empty vector, FLAG-TWIST1 WT or K33R cells were subjected to western blot analysis with the indicated antibodies. (B) RNA extracted from U251 stably expressing empty vector or LINC-PINT cDNA were measured by qPCR. mRNA levels were normalized to GAPDH and then to empty vector cells. Error bars are SEM.

